# Effect of plastic and seaweed shelters on the skin microbiome of lumpfish *Cyclopterus lumpus* used as cleaner fish in aquaculture pens

**DOI:** 10.1101/2025.03.21.644487

**Authors:** Ása Jacobsen, Agnes Mols Mortensen, Kirstin Eliasen, Elin Egholm, Ása Johannesen

**Affiliations:** Department of Biotechnology, Firum PF, Tórshavn, The Faroe Islands; Tari Spf., Fámjin, The Faroe Islands; Department of Fish Health, Firum PF, Hvalvík, The Faroe Islands; Department of Ecology, Firum PF, Hvalvík, The Faroe Islands

**Author notes:** Corresponding author (ÁJ).

## Abstract

Atlantic salmon (*Salmo* salar) aquaculture is a major industry in several countries worldwide and a growing enterprise in others. One of the main challenges the industry faces is infestations with the parasitic copepod *Lepeoptheirus salmonis*, or salmon lice. Several different chemical and mechanical methods are available for alleviating the problem, but often at cost to salmon welfare and/or the environment. In some regions cleaner fish have been introduced to farming facilities as an environmentally and salmon welfare friendly option for reducing the sea lice infestations. In some North Atlantic countries, lumpfish (*Cyclopterus lumpus*) are being used as cleaner fish. However, the welfare and high mortality rates of lumpfish in salmon farming is frequently a problem, and the need to improve lumpfish welfare is great. One adaptation for salmon farms is to provide the lumpfish with shelters to meet their need to rest and hide. Plastic shelters are the most widely used form, but seaweed shelters have more recently also been applied as a more natural solution. This project investigated the potential effect of seaweed and plastic shelters on the skin and gill microbiome of lumpfish and any potential correlation to their welfare. In an experimental setup in a commercial salmon farming facility, lumpfish from pens with either plastic or seaweed shelters were sampled over a period of approximatly three months. The results showed that the bacterial communities on the two shelter types were significantly different and fewer potentially pathogenic bacteria dominated the skin microbiota of lumpfish living with seaweed shelters than of those living with plastic shelters. No differences were detected in the welfare of the lumpfish and further investigations are needed to clarify any potential implications of the differences detected in the skin microbiota of lumpfish including responses to stressful conditions.

## Introduction

Atlantic salmon (*Salmo salar*) aquaculture is a major industry in several North Atlantic countries. Although highly successful, there are many challenges in the industry and one of the main challenges is infestations of salmon lice (*Lepeoptheirus salmonis*) (1), an ectoparasitic copepod that feeds on the mucus, skin and blood of its host. To mitigate this problem various chemical and mechanical solutions have been and are currently being used with various effectiveness. However, there are several drawbacks of these approches such as deminishing effects of medical treatments over time, negative environmental impact and/or substantial salmon welfare issues (2).

In an attempt to avoid these concerns, cleaner fish have been introduced as a biological control to reduce the sealice infestation without adverse effects on salmon welfare or the environment. In the Faroe Islands and Norway, lumpfish (*Cyclopterus lumpus*) are being used as cleaner fish in the aquaculture industry and although this approach has shown to be relatively effective (3–5), lumpfish are a new species in aquaculture and have quite different abilities and requirements than salmon. Therefore, challenges regarding their welfare are concern, as mortality rates are high due to a range of unresolved issues (6,7). A main problem is the high frequency of bacterial infections by known pathogens such as *Tenacibaculum* spp, *Pseudomonas* spp, *Aeromonas* spp. and *Vibrio* spp. (8), which are more likely to occur due to lumpfish already being weakened by other factors (9).

The fish mucosal barriers contain both a range of immunogenic compounds as well microbial communities and in healthy fish the immune system is able to maintain a homeostasis without either infections or excessive immune responses (10,11). At the same time, the composition of the bacterial communities in the mucus is affected by various factors such as the host genetics, environment and diet (10,11). As an example of how the environment affects the fish microbiota, sea anemones have been shown to affect the skin microbiota of fish that live in close contact with them (12). On the other hand, several studies have found that the skin microbiota shows a limited correlation to the bacterial community in the surrounding water (13,14).

Because lumpfish are not adept swimmers and lack a swim bladder, salmon pens with lumpfish are usually equipped with shelters. These are essential for lumpfish welfare as they offer the possibility to hide, rest, and thrive in salmon farming facilities (15). These shelters have usually been constructed of plastic, but in the recent years natural seaweed shelters have gained increasing awareness as a more natural alternative. Seaweeds are well documented to contain a range of antibacterial components (16), which might affect the microbiota of fish living in near contact with them. Here, we describe a study perfomed to investigate the potential effect plastic and seaweed shelters have on the skin microbiota and hence health of lumpfish in aquaculture pens. Since very limited knowledge is available on lumpfish mucosal or skin microbiota the results from this study can provide valuable contributions to the effort of improving lumpfish welfare in aquaculture facilities and to any potential future investigations into the circumstances resulting in detrimental infections. In addition, the knowledge gained can be useful for other fish species used as cleaner fish that require shelters in the aquaculture industry that require shelters.

## Material and methods

### Sampling procedure

The experiment took place in the Faroe Islands (Fig 1a) at a salmon farming facility owned and operated by Bakkafrost. The farming site contained 12 salmon farming pens of which the project had access to sampling and measurements in six, three containing plastic shelters and three containing seaweed shelters (Fig 1b). No lumpfish were added to pens without shelters.

**Fig 1.**
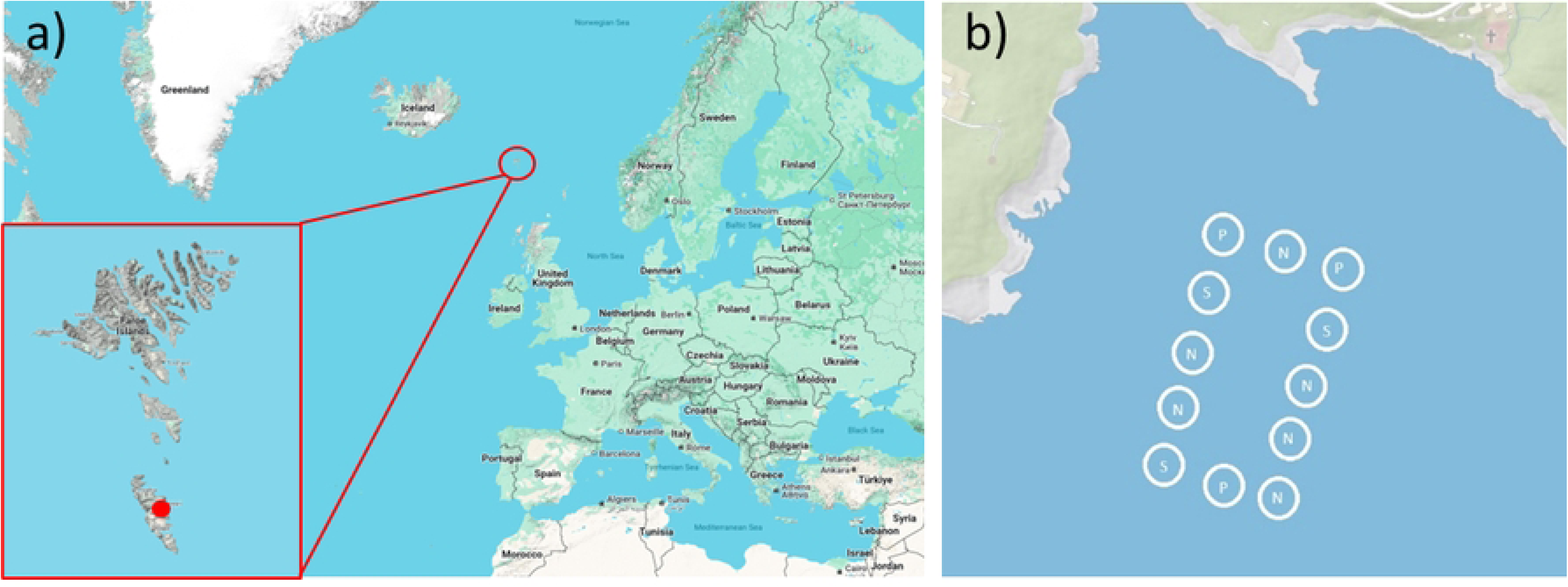
Experimental site. Geographic location of fieldwork in the Northeast Atlantic Ocean, The Faroe Islands (Map data ©2024 Google) (a), and experimental setup in Froðba, The Faroe Islands (Map data ©2024 Google) (b). Salmon farming pens contained seaweed (S), plastic (P), or no (N) shelters.

Lumpfish were added to the pens in April and May 2023 where each pen received 12,500 lumpfish only once but not all pens on the same date. Plastic and seaweed shelters were added to pens prior to deployment of lumpfish. The plastic shelters were of the type SeaNest (Imenco, Norway) with a surface area of 120 m2 per shelter. Two shelters were used in each of the pens with plastic shelters. The seaweed shelters were of the type AkvaNest (Tari, Faroe Islands) and the species used was *Saccharina latissima,* which were propagated onto lines in tanks and grown to appropriate size before deployment in the farming pens. Approximately 60 meters of lines with seaweed was deployed in each pen. Two master lines were stretched across the pen diameter with line droppers with seaweed hanging down from the master line every 5 meters.

From the time of lumpfish deployment and until the end of the experiment in August various monitoring and sampling took place. Every two weeks, 10 lumpfish from each of the six cages were sacrificed and measured using various welfare parameters as described in Eliasen *et al.* (17). Sampling for microbiome analyses was performed of lumpfish skin and gills on three occasions using the same individuals that were used for welfare measurements. Microbiome sampling was performed within ∼2 weeks following deployment and again after ∼8 weeks and ∼14 weeks. In order to allow sampling to follow this strategy, lumpfish from all pens were not sampled at the same date but rather at certain intervals following deployment. Lumpfish retreived from each pen were kept cool and in separate sterile plastic bags until arrival at the field station at the farming site and sampled within an hour. Sampling of skin and gill mucus was performed using sterile swabs prior to welfare measurements. Care was taken to use clean sterile gloves and a sterilised workbench during sampling and sterile plastic tubes for the swabs. The tubes with swabs were immediately stored in dry ice until arrival at the lab where they were stored at –18 °C before DNA extraction within a forthnight. Sampling of the shelters was also performed using swabs. Shelters were raised from the pens and triplicate samples were taken from shelters in each pen at each sampling period. In addition, the shelters were monitored for biofouling each sampling period and categorized from 1 (no biofouling) to 3 (heavy biofouling) in 0.5 increments according to level of biofouling. For seaweed, each sample point was based on 60 measurements of individual leafs, 20 from each of the three pens with seaweed shelters.

Contrary to the seaweed shelters, the plastic shelters were a continuous structure and therefore the scoring was performed for each shelter as a whole, two in each of the three pens with plastic shelters. However, at the last sampling stage the scoring was based on one shelter from each cage due to shortage of manpower at the farming facility. General condition, including tear, holes, and pests for all shelters was estimated by the use of ROV imagery and manual measurements and since catagorized in the same manner as biofouling. In addition, size and growth of the seaweed shelters was measured during sampling. An overview of measurements and sampling performed can be seen in Table 1.

**Table 1.** Overview of sampling and measurements.

| Stage | Date | Sample origin | Samples and measurements |
| --- | --- | --- | --- |
| A | 19.04.2023 | Lumpfish (n=10) pen no. 4,12 | Microbiome gills & skin, welfare |
|  | 03.05.2023 | Lumpfish (n=10) pen no. 1, 3, 7, 8 | Microbiome gills & skin, welfare |
|  | 22.05.2023 | Shelters | Microbiome, biofouling, condition, growth |
| B | 31.05.2023 | Lumpfish (n=10) pen no. 4,12 | Microbiome gills & skin, welfare |
|  | 20.06.2023 | Lumpfish (n=10) pen no. 1, 3, 7, 8 | Microbiome gills & skin, welfare |
|  | 20.06.2023 | Shelters | Microbiome, biofouling, condition, growth |
| C | 10.07.2023 | Lumpfish (n=10) pen no. 4,12 | Microbiome gills & skin, welfare |
|  | 26.07.2023 | Lumpfish (n=10) pen no. 1, 3, 7, 8 | Microbiome gills & skin, welfare |
|  | 07.08.2023 | Shelters | Microbiome, biofouling, condition, growth |

### DNA extraction, library preparation, and sequencing

DNA extraction was performed using the Soil DNA Isolation Plus Kit (Norgen Biotek, Canada). The lysis buffer from the kit was added to the tubes containing the swabs. The samples were since incubated at room temperature for 20 minutes with vortex mixing every 5 minutes. The liquid was then transferred to Bead Tubes from the kit before proceeding with the extraction protocol as described by the manufacturer. Elution volume was 80 ul.

Library preparation was performed using the Quick-16S NGS Library Prep Kit (Zymo Research, Germany) but using the 341f/785r amplicon primers (18,19) and Nextera Index primers (Qiagen, Germany). DNA concentrations were measured using the Qubit dsDNA HS kit (Invitrogen, USA) and Clariostar plate reader (BMG Labtech, Germany). Libraries were sequenced at Novogene (Cambridge, UK) using an Illumina (San Diego, USA) platform producing 2×300 bp paired end reads.

### Data analysis

Raw sequence data were processed using Qiime2 (20) following the methods described in (21). Once taxonomy, alpha diversity, and beta diversity were obtained, feature tables as well as core metric results and taxonomies were imported into R (22) with “tidyverse” (23) for analysis and plotting using the package “qiime2R” (24). Data were further processed and filtered for plots using “metacoder” (25). Unassigned OTUs as well as Eukaryota, Mitochondria, and Chloroplasts were removed for downstream analyses.

In order to compare alpha diversity metrics, mixed effects linear models were constructed using the package “lme4” (26) with “lmertest” (27). First, diversity by sample type was compared with pen as random factor. Then the effect of shelter type was investigated on both diversity on gills and skin again using pen as random factor. Finally, the effect of sample time was investigated using sample type as random effect.

Due to the high diversity, differential abundance analyses were carried out on the top 30 relative most abundant taxa to avoid excessively high multiple correction rates. Differential abundance was tested using “corncob’s” (28) differentialTest function, which runs a model using maximum likelihood for the Beta-binomial distribution with a logit link function for each taxon. P values were adjusted for multiple testing using the built in false discovery rate control function. One test was carried out for all samples comparing those from pens with plastic shelters to those with seaweed shelters and controlling for type of sample, that is what type of substrate the sample was from. Subsequent tests compared relative abundance on shelters, gills, and skin in cages with plastic and seaweed shelters. Finally differential abundance over time was tested while controlling for sample type and whether samples came from pens with seaweed shelters or plastic shelters. The output from the models is presented in figures and tables using the expected difference in the logit-transformed relative abundance between the two groups and 95% confidence intervals.

Beta diversity analysis was based on a Bray-Curtis dissimilarity matrix and illustrated with a PCoA plot. A permanova test using the Adonis2 function was applied to test for potential effects of the variables sample type, sampling stage, and shelter type. Shelter biofouling data were analysed using a clm from the package “ordinal” with shelter type as predictor and score as ordered response.

## Results

### Shelters

The surface area of the seaweed shelters was estimated based on measurements performed at sampling. In May the seaweed leaves were on average 94.6 cm (SD = 23.0) and 8.8 cm (SD = 2.2) in length and width respectively, while in August these values were 137.9 cm (DS = 30.2) and 13.8 cm (SD = 3.1). With approximately 170 individuals per metre of the 60 m line per pen, and adjusting for the shape of the seaweed leaves and non-useable areas by a reduction of 75 % and 50%, respectively, the estimated surface area of the seaweed shelters in each pen was between 637 m^2^ to 1,456 m^2^ from deployment to the end of the experiment. In comparison, the plastic shelters had a surface area of 240 m^2^ per pen.

Based on the biofouling measurements, a cumulative link model showed significant differences between biofouling on the two shelter types (z = −6.151, N = 195, P < 0.001). The plastic shelters had more biofouling than the seaweed shelters at all three measuring periods and had heavy biofouling in all three pens at the last measuring (Fig 2a). The seaweed shelters on the other hand never had more than light biofouling (Fig 2b). Hence, the results from the biofouling measurements suggest that plastic shelters are more prone to biofouling than seaweed shelters, which however can be mitigated by cleaning procedures.

**Fig 2.**
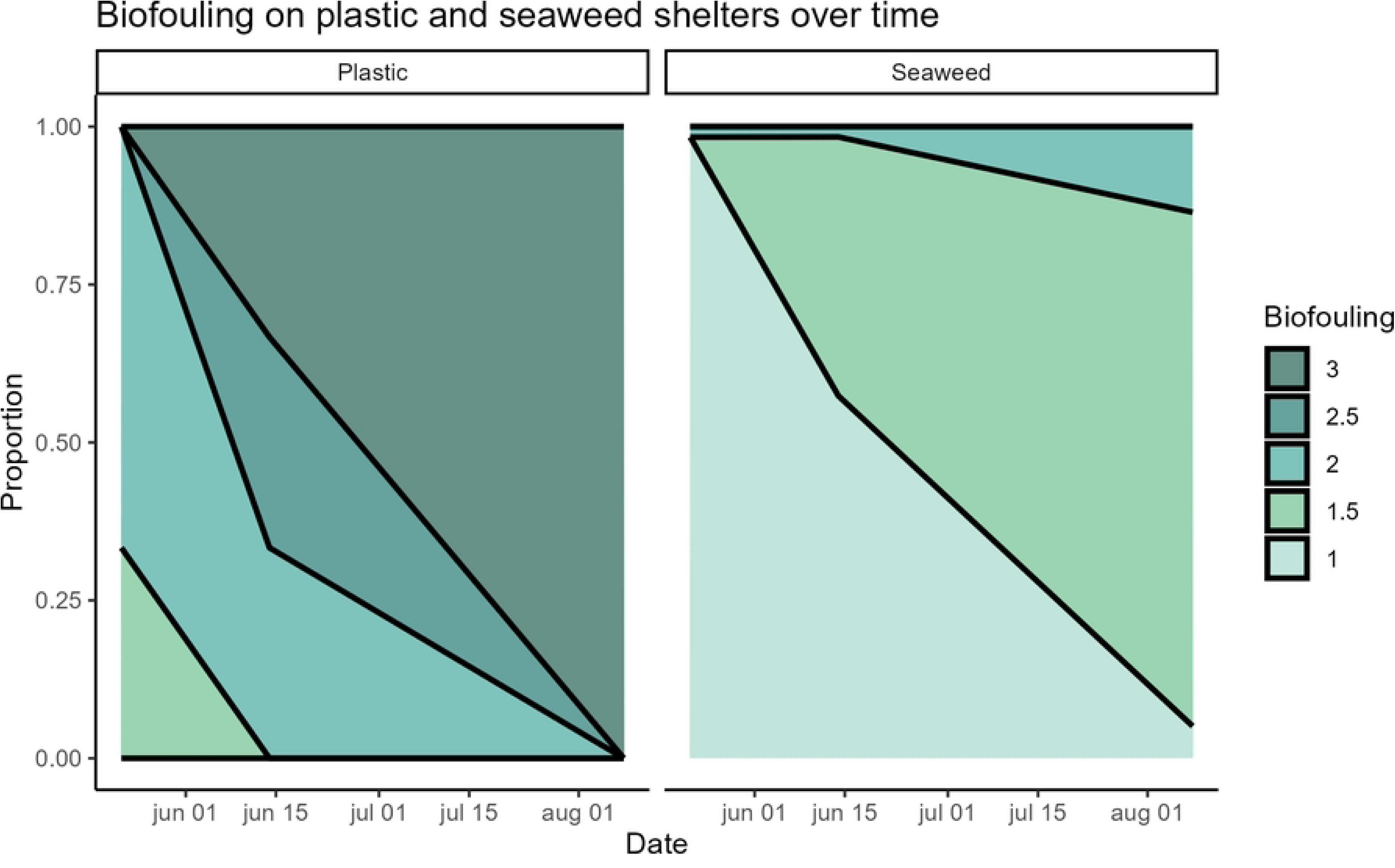
Shelter biofouling. Level of biofouling on seaweed (a) and plastic (b) shelters. Biofouling categories: 1 = no biofouling; 1.5 = sparse biofouling; 2 = full coverage of light biofouling; 2.5 = full coverage of medium biofouling; 3 = full coverage of heavy biofouling. On the other hand, the plastic shelters were in good condition throughout the experiment while the condition of the seaweed shelters deteriorated slightly (data not shown).

### Sequencing data

From the 414 samples sequenced, the total number of reads after quality filtration and removal of ASVs with less than 10 reads was 311,894,277 with a median of 673,995 reads per sample. One seaweed sample only had 1,811 reads and was discarded for downstream analyses while the second lowest value was 34,994 reads.

Four negative control swabs, opened at various times during field sampling, were processed alongside the other samples. DNA concentrations following DNA extraction were either too low to detect or max 0.5 ng/ul. Following library prep where low concentration samples were subject to more PCR cycles, the DNA concentrations were more similar to that of the other samples.

They were since sequenced with the other samples and produced on average 503,066 quality filtered reads. The most dominating bacteria in the negative controls were *Methylobacterium* and *Cutibacterium*. This was taken into account in the downstream data interpretation.

### Alpha diversity

Alpha diversity analysis of the various sample types showed a significant difference in Shannon diversity of the bacterial communities between all sample types (see Table 2 for details).

**Table 2:** Alpha diversity statistics.

| Comparison |  | vs | Shannon entropy | Faith's PD |
| --- | --- | --- | --- | --- |
| Sample types | Gills | Skin | t = 8.07, P < 0.001*** | t = 1.38, P = 0.169 |
|  |  | Plastic | t = 17.92, P < 0.001*** | t = 2.84, P = 0.005** |
|  |  | Seaweed | t = 7.11, P < 0.001*** | t = -0.03, P = 0.979 |
|  | Skin | Plastic | t = 13.92, P < 0.001*** | t = 2.15, P = 0.032* |
|  |  | Seaweed | t = 3.24, P = 0.001** | t = -0.69, P = 0.488 |
|  | Seaweed | Plastic | t = 7.61, P < 0.001*** | t = 2.08, P = 0.038* |
| Gills | Seaweed | Plastic | t = -0.86, P = 0.437 | t = 3.82, P = 0.019* |
| Skin | Seaweed | Plastic | t = -1.55, P = 0.196 | t = -1.33, P = 0.254 |
| Time | A | B | t = 5.29, P < 0.001*** | t = -0.51, P = 0.608 |
|  |  | C | t = 4.94, P < 0.001*** | t = 4.48, P < 0.001*** |
|  | B | C | t = -0.33, P = 0.740 | t = 4.99, P < 0.001*** |

Linear mixed effects models comparing alpha diversity metrics (Shannon entropy and Faith pd) over sample type (tissue type and shelter), gills and skin over shelter availability in pen, and samples over time. The t and P values are from the summary tables from the models, which provide differences between each factor level. Significance levels: \* p < 0.05, \*\* p < 0.01, \*\*\* p < 0.001

Plastic shelters had the highest Shannon diversity while gills had the lowest (Fig 3a). Regarding the phylogenetic diversity, only plastic shelters differed significantly from the other sample types (Table 2), having the highest Faith’s PD values (Fig 3b).

**Fig 3.**
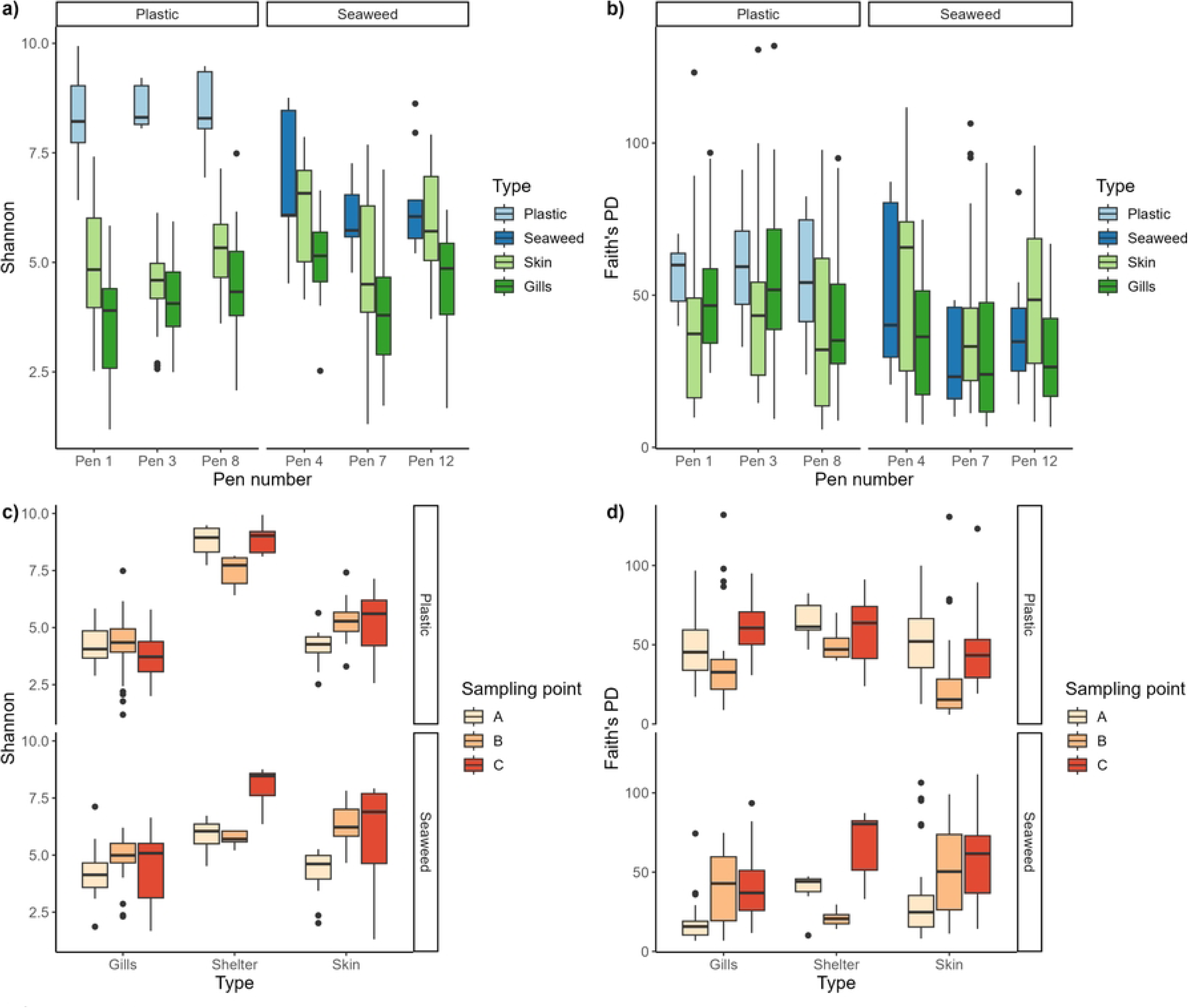
Alpha diversity. Alpha diversity measures Shannon (a) and Faith’s PD (b) for all sample types in each pen and Shannon (c) and Faith’s PD (d) alpha diversity measures for the various sample types over time in pens with plastic and seaweed shelters.

The microbial community on gills of lumpfish living in pens with plastic shelters also had a significantly higher phylogenetic diversity than those on gills of lumpfish living in pens with seaweed shelters (t = 3.82; p = 0.019, Fig 3b). In pens with seaweed shelters there was a general trend of increasing shannon and phylogenetic diversity over the sampling period for all sample types while in pens with plastic shelters the diversity values were more fluctuating (Fig 3cd). The phylogenetic diversity at the last sampling period was significantly different than of the previous two sampling periods, and there was also a significant difference in Shannon diversity between sampling stage A and C (see Table 2). This indicates changes in the microbiota over time although not necessarily the same changes in pens with seaweed and plastic shelters. An overview of the alpha diversity statistical analysis and results can be seen in Table 2.

### Composition of bacterial communities

Proteobacteria was the dominating phyla in all sample types (Fig 4). However, Alphaproteobacteria had higher relative abundance than gammaproteobacteria in both shelter types while it was the opposite for lumpfish skin and gills. On plastic shelters, Rhodobacterales was the single most dominating order of the Alphaproteobacteria (Fig 4a) while Caulobacterales were equally dominating on seaweed shelters (Fig 4b). On skin and gills there were several prominent Alphaproteobacteria orders although Rhodobacterales also was the most dominant order there (Fig 4cd).

**Fig 4.**
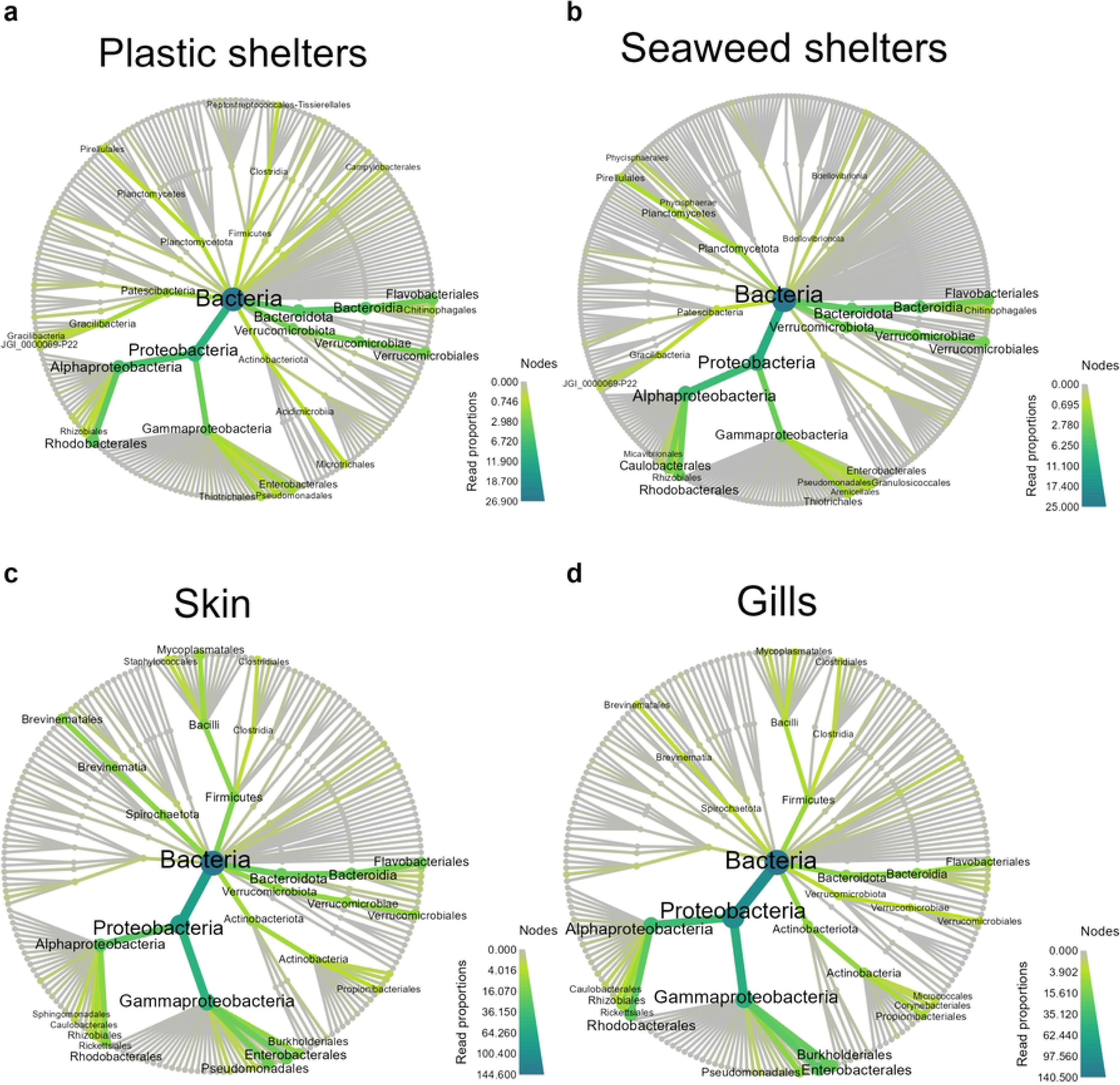
Heat trees. Phylogenetic heat trees down to order level for microbiome on plastic (a) and seaweed (b) shelters, and lumpfish skin (c) and gills (d) The orders in Gammaproteobcateria with the highest relative abundance in the lumpfish samples were Enterobacterales, Pseudomonadales, and Burkholderiales (Fig 4cd). Burkholderiales was not prominent on either shelter type but Thiotrichales was relatively abundant (Fig 4ab).

Bacteroidota and Verrucomicrobia were also prominent phyla on the shelters, but less so in the lumpfish samples. Other phyla detected were Planctomycetota and Patescibacteria, mainly in seaweed and plastic shelters respectively (Fig 4ab), Firmicutes and Spirochaetota, mainly in skin samples (fig 4c), and Actinobacteriota in gill samples (fig 4d).

Almost half of the bacteria detected on the shelters were exclusively found on plastic (25.7 %) or seaweed (18.8 %) shelters, indicating quite different microbial communities on the two shelter types. The differences between plastic and seaweed shelters, illustrated in the bacterial community as a whole, were also apparent when looking at the most dominant bacteria, as there were not the same on seaweed and plastic shelters. Of the 17 most relative abundant genera depicted on the barplots (Fig 5ab) only eight were on both shelter types.

The top 17 dominating bacterial genera on the plastic shelters consisted of a lower percentage of the total reads than those on seaweed shelters, aligning with the higher Shannon diversity detected in the plastic samples. Among the dominating bacteria found on both shelter types were mostly well known marine genera such as *Polaribacter*, *Sulfitobacter*, and *Colwellia* that are widely distributed, especially in the colder regions (29–31). *Rubritalea* was the most dominating genus on plastic shelters and *Fusibacter*, which is mostly known for containing various pathogens affecting humans and some terrestrial animals, was also among the prominent bacteria on plastic shelters. Recently, however, a study detected an aero-, halo-, and psychrotolerant *Fusibacter* strain inhabiting the marine sediment in an arctic region, suggesting that *Fusibacter* could be more widely distributed than previously thought (32).

Overall, there were similar patterns of dominating bacteria on the same shelter types in the various pens, but some changes were apparent over time, especially on plastic shelters (Fig 5ab). Statistical comparisons of the bacterial community compositions on the two shelter types showed a significant difference in the majority of the 30 genera with highest relative abundance (Fig 5c). Among these genera was the pathogenic genus *Tenacibaculum,* which was the second most abundant genus detected on plastic shelters but was not identified among the dominant bacteria on seaweed shelters (Fig 5ab). Among the genera with significantly higher relative abundances on seaweed shelters were *Dokdonia* and *Blastopirellula,* previously detected as part of the biofilm forming bacterial communities on various macroalgae (33–35).

**Fig 5.**
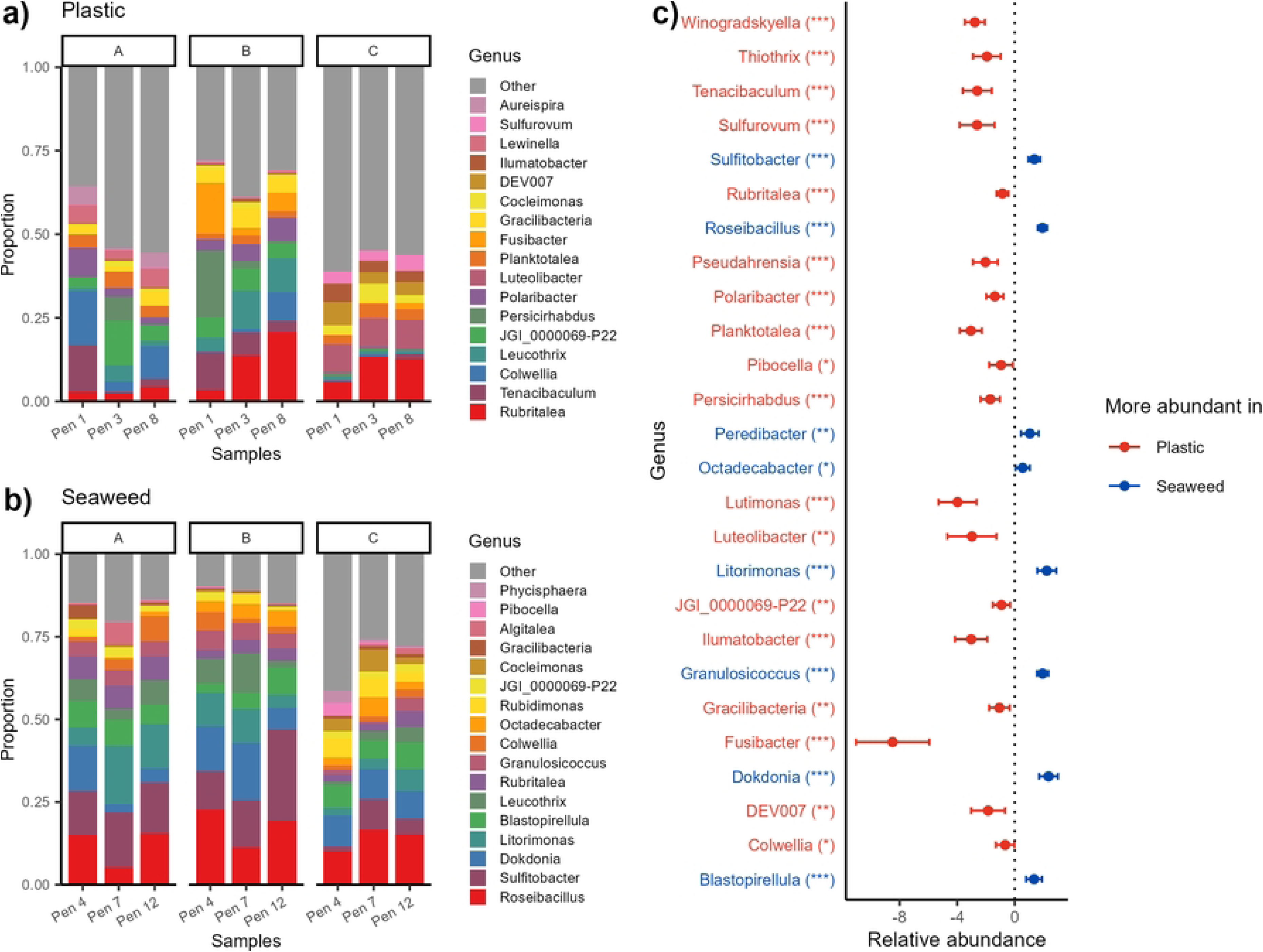
Taxonomic composition of bacterial community in shelter biofilm. Bacterial community compositions illustrating the most dominating genera on plastic (a) and seaweed (b) shelters split by pen (x axis) and sampling point (panel A to C). Differences in relative abundance of the top 30 most prevalent genera on shelters found to be significantly different depending on shelter type (c). Red genera (to the left) are more abundant in plastic shelters while blue genera (to the right) are more abundant on seaweed shelters. Significance levels: * p < 0.05, ** p < 0.01, *** p < 0.001.

Analysis of the lumpfish skin microbiome showed a change over time in dominance of the major groups of the bacterial community. This shift was apparent in the skin of lumpfish living in pens with both plastic and seaweed shelters (Fig 6ab). The lumpfish from the triplicate pens with same shelter type also seemed to have more similar skin microbiota early after deployment and differ more at the other two sampling points (Fig. 6ab). In the top 30 most relatively abundant genera, fourteen genera differed significantly in abundance between the skin of lumpfish in pens with seaweed shelters and those with plastic shelters (fig 6c).

**Fig 6.**
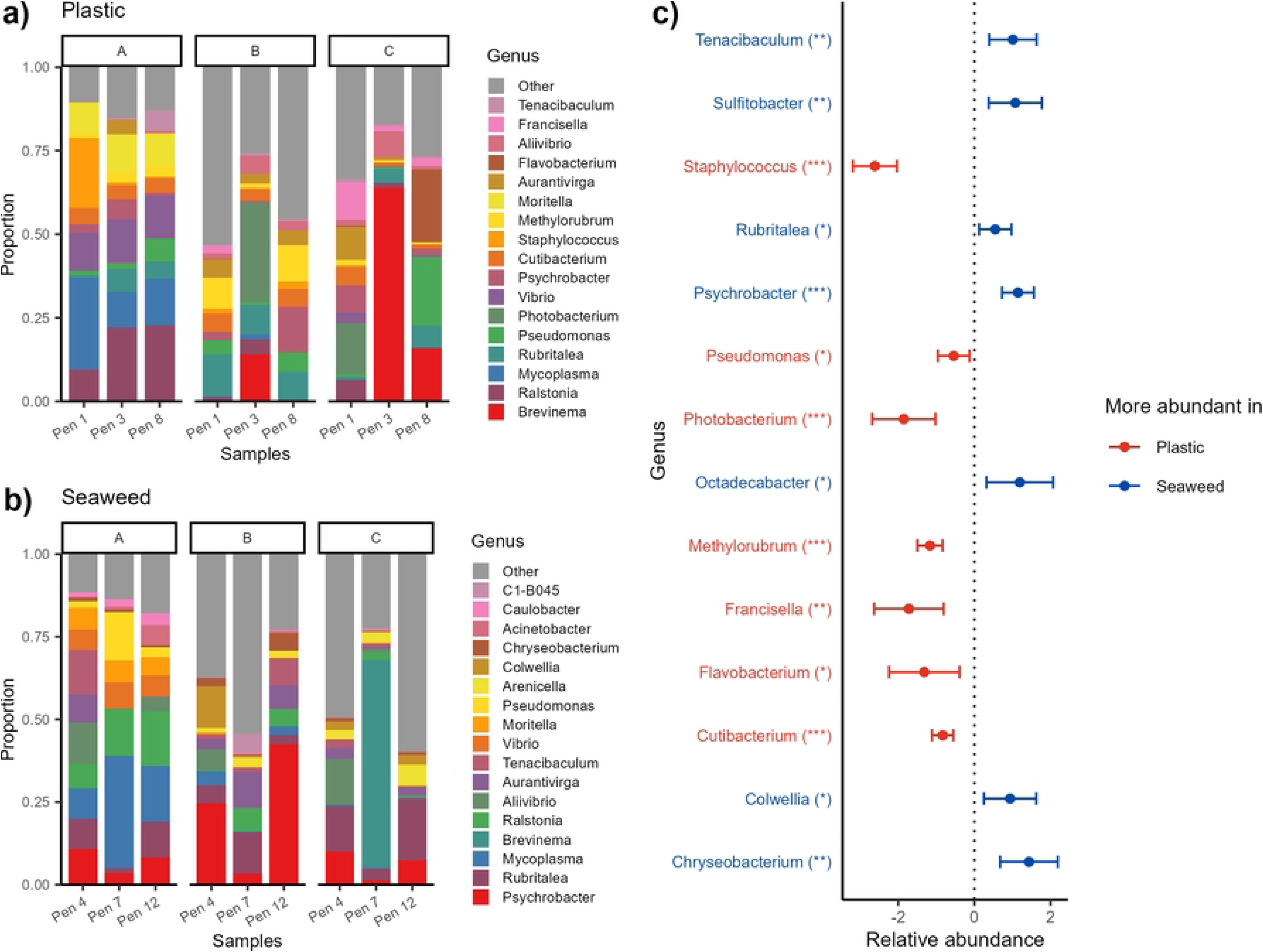
Taxonomic composition of bacterial community in lumpfish skin mucus. Bacterial community compositions illustrating the most dominating genera on lumpfish skin from pens with plastic (a) and seaweed (b) shelters split by pen (x axis) and sampling point (panel A to C). Differences in relative abundance of the top 30 most prevalent genera on shelters found to be significantly different depending on shelter type (c). Red genera (to the left) are more abundant in plastic shelters while blue genera (to the right) are more abundant on seaweed shelters. Significance levels: * p < 0.05, ** p < 0.01, *** p < 0.001.

The eleven relatively most abundant genera in the skin microbiota of lumpfish from pens with seaweed shelters were also among the 17 genera of highest relative abundance on lumpfish from pens with plastic shelters (Fig. 6ab). The genera *Tenacibaculum*, *Vibrio, Moritella, Pseudomonas* and *Aliivibrio* with known pathogenic species (36,37) were detected among the most dominant genera on lumpfish from pens with both shelter types. However, *Tenacibaculum, Moritella* and *Vibrio* were mainly present shortly after the lumpfish were deployed and were much less dominant at the later two sampling periods (Fig 6ab). In addition to these, several other potential pathogens such as *Francisella* (38)*, Staphylococcus* (39)*, Flavobacterium* (40) were also detected among the dominant genera in skin samples from lumpfish living in pens with plastic shelters (Fig 6b). These four potentially harmful genera all had significantly higher relative abundance on lumpfish from pens with plastic shelters, while *Tenacibaculum* was significantly more abundant on lumpfish from pens with seaweed shelters (Fig. 6c). However, the potentially beneficial bacteria *Psychrobacter* (41) was the most dominant genus in skin samples from lumpfish living in pens with seaweed shelters. In contrast, lumpfish living in pens with plastic shelters had significantly lower relative abundance of *Psychrobacter* in the skin microbiota. At the last sampling period, the genus *Brevinema* was also highly noticable in the skin samples of lumpfish from pens 7 and 3 containing seaweed and plastic shelters respectively.

Many of the dominant genera detected in the skin samples were also among the dominant genera in the gills samples (Fig. 7ab). *Ralstonia*, which was among the dominating genera in skin samples, was the genus with highest relative abundance in gills of lumpfish living with both shelter types.

Several of the dominating genera detected in lumpfish skin (Fig 6ab) were also among the dominating genera in lumpfish gills (Fig 7ab). In addition, twelve of the seventeen most dominating genera in the gills microbiota of lumpfish in pens with seaweed and plastic shelters were the same (Fig 7ab), similar to the pattern seen with skin sample s. However, the gill microbiome of lumpfish living in pens with seaweed and plastic shelters was more similar than what was the case for the skin microbiota, as only four genera among the top 30 most dominating genera were significantly different (Fig 7c). The only genus of the four with significantly different relative abundance that was more prominent in lumpfish living in pens with seaweed shelters was *Psychrobacter.* On the other hand, one of the three genera with significantly higher relative abundance in skin of lumpfish living in pens with plastic shelters was *Photobacterium*, which contain species that can be pathogenic in warmer waters (42). Other dominating genera were well known marine bacteria such as *Colwellia*, which is widespread in the colder marine regions (43), *Cutibacterium* commonly found in the gut microbiota and sometimes gills of marine fish (44,45) and *Mycoplasma* that often is a dominant part of the salmon gut microbiota (46,47).

**Fig 7.**
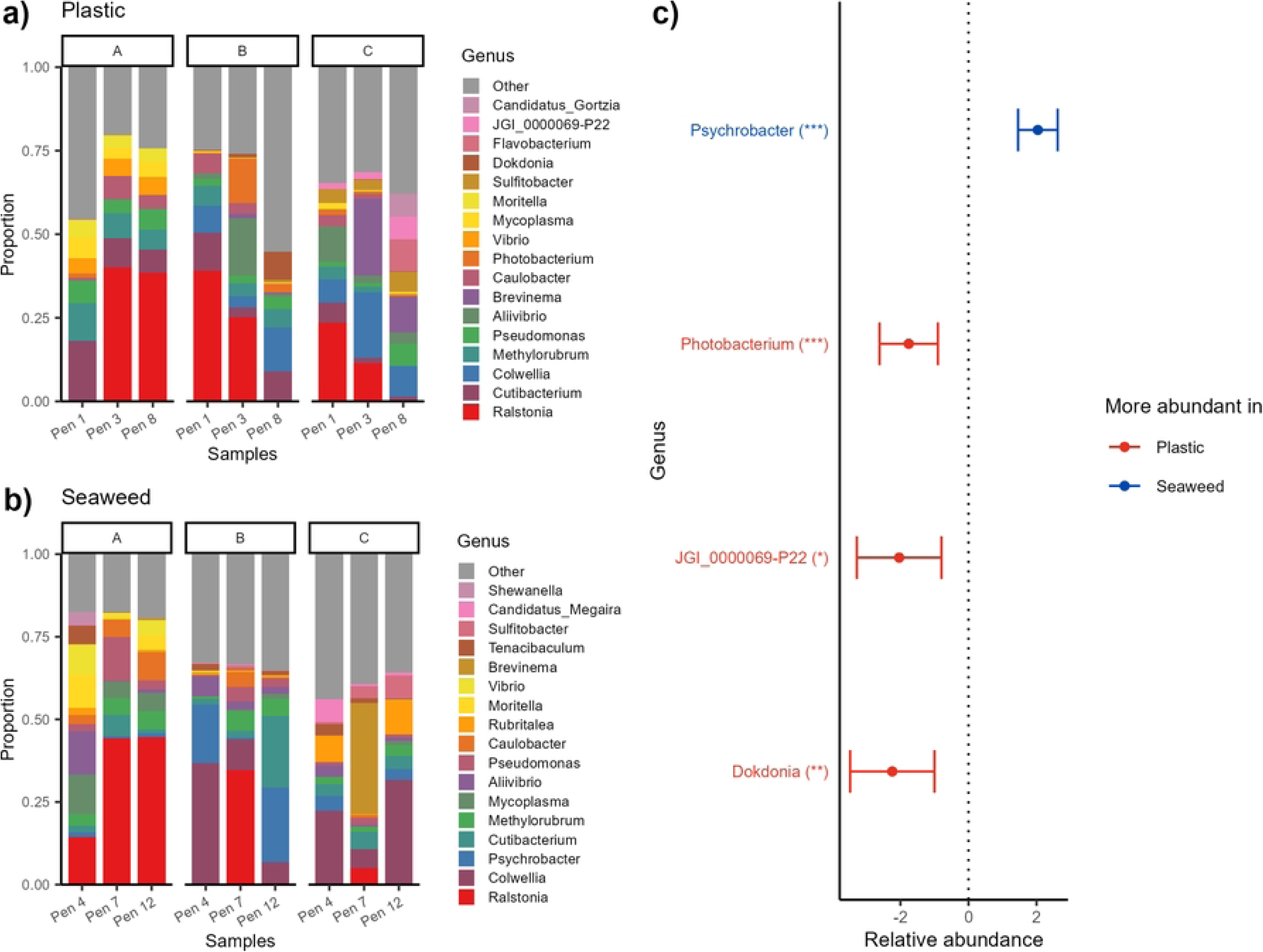
Taxonomic composition of bacterial community in lumpfish gill muscus. Bacterial composition of the most dominating genera in lumpfish gills in seaweed (a) and plastic (b) shelters split by pen (x axis) and sampling point (panel A to C). Differences in relative abundance of the top 30 most prevalent genera on lumpfish gills found to be significantly different depending on shelter type. Red genera (to the left) are more abundant in lumpfish living with plastic shelters while blue genera (to the right) are more abundant in those living with seaweed shelters. Significance levels: * p < 0.05, ** p < 0.01, *** p < 0.001.

### Beta diversity

The beta diversity analysis, illustrated in a Bray Curtis PCoA plot (Fig 8), showed a change occurring over the experimental period. Skin and gill microbiomes in yellow and purple, respectively, were partially intertwined and/or clustered in close proximity to each other early after deployment. Over time the variation described changed from PC1 to PC2 and the two sample types had a little less overlap. On the other hand, the microbiota on the shelters was separated from the lumpfish samples at the first sampling stage, but over time the skin microbiota separated somewhat from the gill samples and drew closer to the shelter samples. The gill microbiome seemed less affected by the shelters as the gill samples were distanded further away from the shelter.

**Fig 8.**
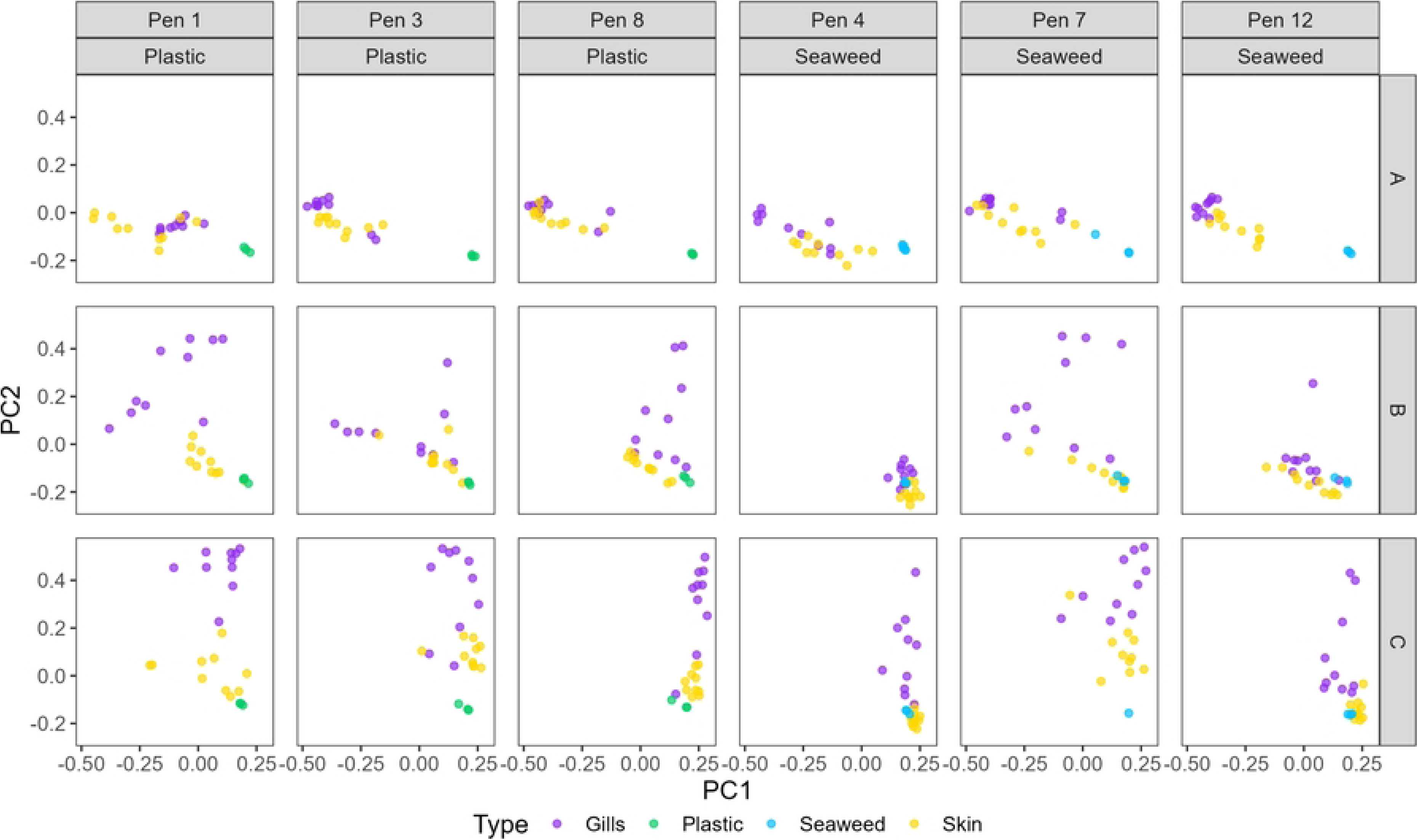
Bray Curtis PCoA. PCoA plot based on Bray Curtis beta diversity distance matrix. Samples are split into columns by the various pens, with those containing plastic shelters to the left and those with seaweed shelters to the right. The rows are the three sampling stages A, B, and C.

A permanova test investigating potential effects of sample type, sampling stage, and shelter type on the Bray Curtis distance matrix was performed. All three parameters had a significant impact on distances between samples (sample type: F_2,409_ = 11.873, p < 0.001; sampling stage: F_2,409_ = 11.518, p < 0.001; shelter type: F_1,409_ = 2.631, p < 0.001). The statistical analysis support what the PCoA plot illustrated (Fig 8), that there was a more visible difference between the sample types and sampling stages than between skin and gills from lumpfish living in pens with either plastic or seaweed shelters. However, there still was a significant difference between the microbiota of lumpfish living with the two shelter types.

### Welfare indicators

The results from the welfare measurements including fin and skin condition, characterized 1 to 3 representing good to bad, showed overall low indicator values for the skin (Fig 9a), meaning the lumpfish were in good condition, while the fin condition was generally higher (Fig 9b).

**Figure 9.**
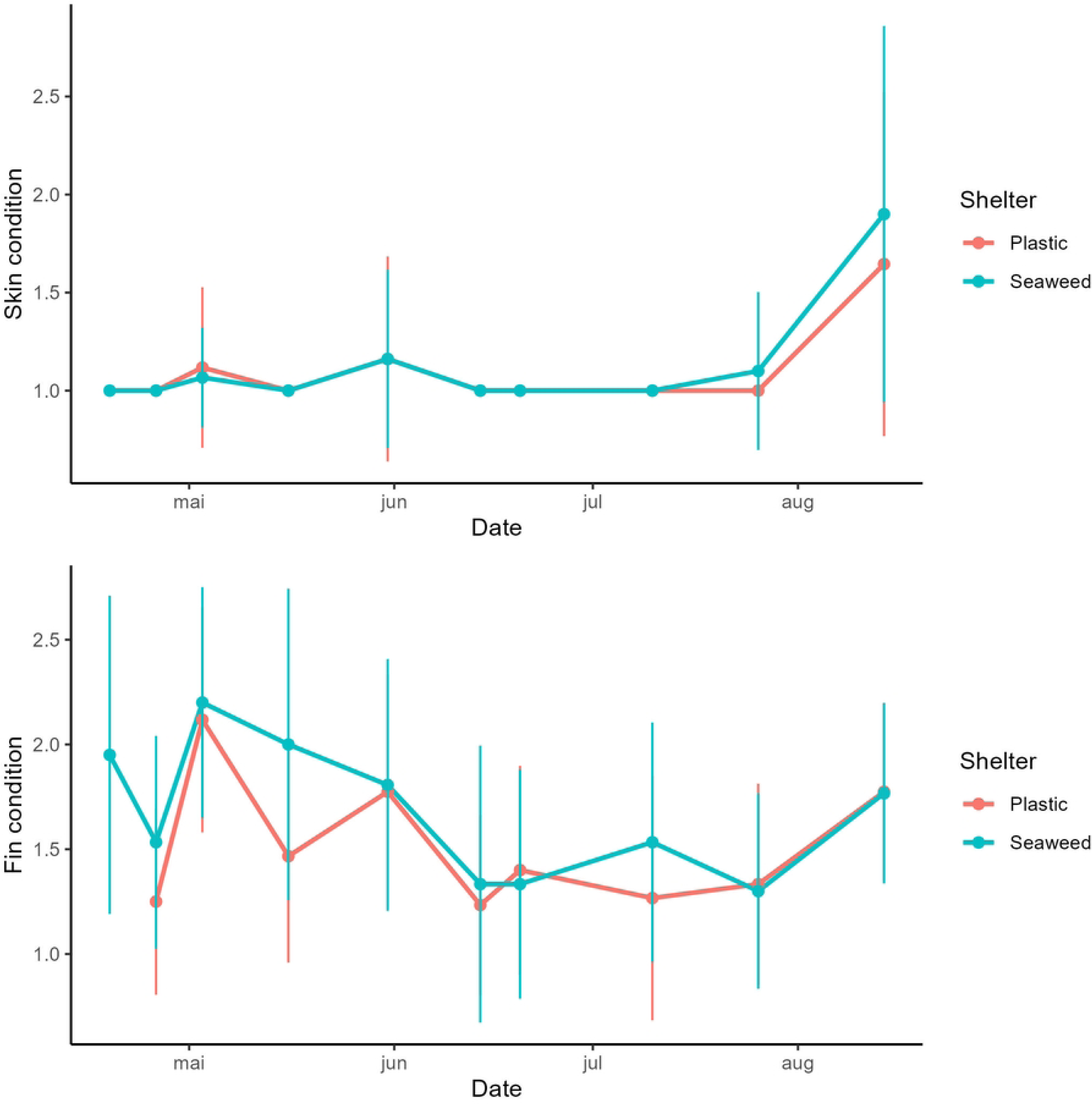
Welfare indicators. Measurements of lumpfish skin (a) and fin condition. The graphs show average and standard deviation of possible categories 1, 2, and 3 representing good medium and bad condition, respectively.

No significant differences in fin and skin condition were detected between lumpfish in pens with plastic and seaweed shelters. Hence, any differences in relative load of potentially harmful bacteria in the skin mucus were not reflected in the welfare indicators of the lumpfish at the time of sampling.

## Discussion

It is critical for the welfare of lumpfish used as cleaner fish in the aquaculture industry that shelters are placed in the pens (48), and that these shelters are suitable and do not cause unintended harm. The plastic shelters investigated in this project were more prone to biofouling than the seaweed shelters used. This in itself suggested that the seaweed shelters were a better environment for the lumpfish that were in regular close contact with the shelters. However, biofouling on the plastic shelters can be mitigated with cleaning procedures, although it might cause some disturbance for the lumpfish, and the plastic shelters did not show any sign of wear and tear in the experimental time frame, while the seaweed shelters did show some signs of degradation. The biofilm microbiota of both plastic and seaweed shelters had significantly higher Shannon diversities than the lumpfish skin and gills microbiota, but the Shannon diversity of the plastic shelter microbiota was also significantly higher than that of the seaweed shelters. The plastic shelter microbiota also had higher phylogenetic diversity than that of the seaweed shelters, indicating that fewer bacterial taxa could thrive on live *Saccharina latissima* leaves, used as seaweed shelter in this case, than on the plastic shelters. Another sign of the difference between the shelter types was that only about half of the bacterial genera detected on the shelters were on both shelter types while the others were solely found on either plastic or seaweed shelters. This was reflected in the high amount of significant differences in the relative content of various bacteria. Among these, a significant difference was detected between the relative abundances of *Tenacibaculum* on the two shelter types. However, the relative abundance of *Tenacibaculum* diminished over time and this pattern was also observed in some of the lumpfish skin samples, suggesting that these bacteria were already part of the lumpfish microbiota when transported to the aquaculture farming site. Earlier investigations have also detected high loads of *Tenacibaculum* in the transport water and in samples from lumpfish being shipped from abroad to the Faroe Islands for use as cleanerfish in the aquaculture industry (pers. com. Esbern J. Patursson). Although the relative abundance of *Tenacibaculum* diminished over time on the plastic shelters, it might be noteworthy that they were never among the dominating genera on the seaweed shelters. This finding is consistent with the results from a study by Liu *et al*. (49) investigating biofilm on cultured *Saccharina latissima* where they detected a much lower relative abundance of *Tenacibaculum* on the seaweed than in the surrounding seawater. Others have also detected a relatively high load of potentially harmful bacteria on plastic in marine aquaculture operations, while the bacterial composition also differs from the surrounding seawater (50,51).

Although no seawater sample was included in this study, the aquaculture site chosen for the experiment was placed in an area with good water exchange providing the best possibility for similar conditions for all pens. The most dominating genus on plastic shelters was *Rubritalea*, consistent with the findings of Papale *et al*. (52) who consistently found *Rubritalea* in biofilm on plastic submerged in seawater. Other genera with significantly higher relative abundance on plastic shelters included the marine genus *Planktotalea* containing only a few documented species detected in colder marine regions (53,54), but noticeably detected in plastic biofilm and not the surrounding seawater in the study by Papale *et al*. (52). Overall, the data indicated that the two shelter types provided different environments for the lumpfish in this study.

The gills of lumpfish living with either shelter type had an overall more similar microbiota than the other sample types. This suggested a higher reflection of the surrounding seawater, which in a farming facility with good water exchange, such as Froðba (55), should have a relatively homogenous mixture of the microbial community coming through the various pens. Other studies have also found that gills had the highest similarity with seawater although there was tissue specific microbial communities on fish (45). Many of the dominating genera present in the gills were also well known ubiquitous marine bacteria or fish gut bacteria. However, we find it interesting that of the 30 most dominating genera only *Psychrobacter* was significantly more relative abundant in gills of lumpfish from pens with seaweed shelters. *Psychrobacter* is often used as a probiotics and has been demonstrated to reduce the negative effects of *Tenacibaculum* in fish (41). *Psychrobacter* was also the genus of highest relative abundance in skin samples from lumpfish living with seaweed shelters and their relative abundance in the skin from lumpfish living in pens with plastic shelters was significantly lower. This might suggest a beneficial effect from the seaweed shelters, although *Psychrobacter* was not detected among the dominating bacteria on the seaweed itself.

The skin samples contained overall many potentially harmful bacteria among the dominating genera. Lumpfish from pens with plastic shelters had a higher number of these taxa with a combined higher relative abundance. In addition, for many of the dominating and potentially harmful taxa detected on lumpfish from pens with seaweed shelters the relative abundance was high shortly after deployment and dimished throughout the sampling period. Only *Aliivibrio* was relatively abundant in one of the pens with seaweed shelters at the final sampling.

In pens with plastic shelters, there was also a shift in microbial composition on the lumpfish skin. *Brevinema*, which has been associated with the gut microbiota of several fish species, including salmon (56), had a high relative abundance in pens with both shelter types at the last sampling.

The presence of *Brevinema* on the lumpfish skin is likely a reflection of the salmon feces in the water and the genus has not been reported to include pathogens. On the other hand, there were still several potentially harmful bacteria among the dominant genera in the lumpfish skin in pens with plastic shelters at the final sampling period e.g. *Pseudomonas, Francisella, Aliivibrio and Flavobacterium*. Combined, these observations suggested that lumpfish living with seaweed shelters had a healthier skin microbiota. However, the welfare measurements did not indicate any differences between lumpfish living in pens with plastic or seaweed shelters. The lumpfish were generally in good condition and it can only be speculated whether more stressful conditions would have resulted in significantly lower welfare for lumpfish in pens with plastic shelters than with seaweed shelters. Also, there is evidence from the literature that the commensal microbiota on fish is species specific (57,58) and that commensal bacteria can also become oppertunistic pathogens (59). Together, these traits make it difficult to determine precisely which bacteria are harmful, commensal or perhaps beneficial at any given time for a specific fish species. However, any condition that might cause a clear dominance of potentially harmful bacteria in the skin mucus of lumpfish used as cleaner fish should be kept in mind as other factors weakening the fish might reveal correlations not previously detected.

The beta diversity analysis showed that over time the skin microbiota of the lumpfish and the shelters became more similar while the gills became more separated from the skin samples and to a lesser extent drew closer to the shelters. The most prominent genera in the negative controls were among the most relative abundant genera in the gill samples from lumpfish living with both shelter types. Since these genera are widely distributed in the marine environment, it could be misleading to disregard these genera due to their presence in the negative controls. We consider the eDNA detected in the negative controls to most likely be an effect of the miniscule amounts of saltwater in the air where negative control tubes were opened during fieldwork. Although the joint water environment likely had some influence on the increased similarity between the lumpfish and shelter microbiota over time, there was a strong indication that especially the skin microbiota was affected by the shelter the lumpfish was living in, correlating with the significant difference between microbial compositions on the skin of lumpfish living in plastic shelters and those living in seaweed shelters.

## Conclusions

The results from this study indicated that plastic and seaweed shelters provided different environments for lumpfish residing amongs them and affected their skin microbiota. Furthermore, the skin microbiota of lumpfish living in pens with plastic shelters contained more potentially harmful bacteria among the dominating genera than the skin microbiota of lumpfish living in pens with seaweed shelters. However, no significant difference was detected in the welfare indicators of the lumpfish, which generally were in good condition. Further investigations are needed to determine if having a skin microbiota more dominated by potentially harmful bacteria is of any significance under more stressful conditions.

## Acknowledgements

We wish to thank Bakkafrost for good collaboration and the staff at the farming facility where the experiment took place for operational assistance. We also wish to thank members of the project reference group, Eskil Bendiksen, Esbern Patursson, Kjetil Heggen, and Marner Nolsøe, for constructive discussions.

## Notes

### Competing Interest Statement

I have read the journal's policy and the authors of this manuscript have the following competing interests: Co-author AMM is co-owner of Tari Spf. that produces Akvanest seaweed shelters used in this study. However, AMM did not take part in or influence molecular sampling strategy decisions, lab work, or data analysis.

